# Optogenetic dissection of Rac1 and Cdc42 gradient shaping

**DOI:** 10.1101/317586

**Authors:** S. De Beco, K. Vaidžiulytė, J. Manzi, F. Dalier, F. di Federico, G. Cornilleau, M. Dahan, M. Coppey

## Abstract

During migration, cells present a polarized activity that is aligned with the direction of motion. This cell polarity is established by an internal molecular circuitry, without the requirement of extracellular cues. At the heart of this circuitry, Rho GTPases spontaneously form spatial gradients that define the front and back of migrating cells. At the front of the cell, active Cdc42 forms a steep gradient whereas active Rac1 forms a more extended pattern peaking a few microns away from the cell tip. What are the mechanisms shaping these gradients, and what is the functional role of the shape of these gradients? Combining optogenetics and cell micopatterning, we show that Cdc42 and Rac1 gradients are set by spatial patterns of activators and deactivators and not directly by advection or diffusion mechanisms. Cdc42 simply follows the distribution of GEFs thanks to a uniform GAP activity, whereas Rac1 shaping requires the activity of an additional GAP, β2-chimaerin, which is sharply localized at the tip of the cell. We find that β2-chimaerin recruitment depends on feedbacks from Cdc42 and Rac1. Functionally, the extent -neither the slope nor the amplitude- of RhoGTPases gradients governs cell migration. A Cdc42 gradient with a short spatial extent is required to maximize directionality during cell migration while an extended Rac1 gradient controls the speed of the cell.

## Introduction

Cell migration plays a major role in various biological functions, including embryonic development, immune response, wound closure and cancer invasion. Cells, either isolated or in cohesive groups, are able to respond to many types of spatially distributed environmental cues, including gradients of chemoattractants^1,2^, of tissue stiffness (durotaxis)^3–5^, and of adhesion (haptotaxis)^6,7^. To sense and orient their migration accordingly, cells need to integrate complex and noisy signals and to polarize along the selected direction. A simple explanation to account for such directed migration would be to consider that external gradients are directly translated into internal gradients. However, recent works^8–10^ point to a two-tiered mechanism. First, a set of signaling proteins (RhoGTPases and Ras) behave as an excitable system that spontaneously establish intracellular membrane-bound gradients, conferring the ability of cells to polarize even in the absence of external stimuli. Second, a sensing machinery based on membrane receptors align the polarization axis along the direction of external gradient cues. In the present work, we address the mechanisms shaping the Rho GTPases gradients at the front of randomly migrating cells.

RhoGTPases are known to play a key role in orchestrating the spatially segregated activities that define the polarity axis of migrating cells. At the cell front, membrane protrusions fueled by actin polymerization push the cell forward, while retraction of the cell back depends on acto-myosin contractility^11–13^. The schematic view is that front-to-back gradients of Cdc42 and Rac1 define the cellular front, while RhoA is mostly active at the back. Cdc42 is known to be required for filopodia formation, through N-WASP-mediated activation of the ARP2/3 complex as well as F-actin bundling proteins such as fascin and formin^11,14^. Conversely, Rac1 is involved in branched actin polymerization and lamellipodia formation, through WAVE-mediated activation of the ARP2/3 complex^15^. RhoA is responsible for stress fiber formation and retraction of the cellular tail through Rho kinase-mediated contraction of myosin II^16,17^. In reality the situation is more complex since RhoA is also active at the very front of migrating Mouse Embryonic Fibroblats^18,19^ and is involved in actin polymerization through Diaphanous-related formins as well as focal adhesions^20,21^. In addition, the Rho GTPase family contains more than the 3 members mentioned, with more than 20 proteins having been discovered^20,22^. Despite the fact that these other members are classified in the three Cdc42, Rac1 and RhoA sub-families, they present overlapping activities.

Three main classes of proteins regulate Rho GTPase activity. Guanine Exchange Factors (GEFs) activate Rho GTPases by promoting the exchange from GDP to GTP, whereas GTPase activating proteins (GAPs) inhibit RhoGTPases by catalyzing the hydrolysis of GTP^23^. A multitude of GEFs and GAPs ensure signaling specificity and fine-tuned regulation. In addition, Guanine-nucleotide dissociation inhibitors (GDIs) are negative regulators of RhoGTPases, extracting them from the plasma membrane and blocking their interactions with GEFs^24,25^. GEFs and GAPs can be localized and activated by upstream factors such as Receptor Tyrosine Kinases or interaction with lipids such as PIP3^26,27^, hereby connecting the polarization machinery with the sensing one. Some of the GEFs, GAPs and GDIs are specific to one single GTPase, but many can activate multiple targets, or even behave as a GEF for a GTPase and as a GAP for anther one^28,29^. In addition, GEFs and GAPs can be activated or inhibited downstream of Rho GTPases^30,31^. Complex crosstalks thus connect Rho GTPases and their interactors, resulting in a signaling network that finely regulates Rho GTPases activities. Although many molecular interactions defining this signaling network have been characterized, we currently have little insight on how these interactions are orchestrated in space to shape Rho GTPase activity patterns in migrating cells.

Positive feedbacks acting on Rac1, Cdc42 and RhoA have been proposed to account for their ability to form gradients spontaneously. Rho GTPase activity pulses would be generated thanks to an excitable system^9^ and specific activators like GEFs would orient and stabilize them^8,32^. Yet, activity patterns governed by excitable systems have a propensity to propagate through the whole cell, and inhibitory mechanisms are required to limit their expansion. Rho GTPases are inefficient enzymes, with a slow GTP hydrolysis rate in vitro^33^ such that additional processes are required to prevent a propagation of active Rho GTPases species by diffusion or actin-driven advection^34–36^, where the term advection refers to the oriented (ballistic) transport of material. Three classes of mechanisms could confine Rho GTPases activities. First, GAPs hydrolyze active Rho GTPases and can shorten their lifetimes in the GTP-bound state over orders of magnitude. Rho GTPase cycles can then be locally regulated by GEF and GAP concentrations, whose distributions along the cell would shape Rho GTPase intracellular gradients^37–39,36^. Second, anchoring or trapping in the cortical acto-myosin network can decrease diffusion considerably. Dense actin meshes are thought to behave as signaling scaffolds that concentrate activators of Rho GTPases. Since Rho GTPases trigger actin polymerization and branching, this mechanism could act as a negative feedback restricting their activity zones. Third, Rho GTPase extraction from the plasma membrane by GDIs can be locally regulated^25^, such that deactivation regions could be set by the activity of GDIs. It is unclear which of these mechanisms (fast cycling, anchoring, or extraction from the plasma membrane) or which combination of them, is responsible for the formation of Rho GTPase intracellular spatial patterns.

In this work, we address the mechanisms setting the spatial properties of Cdc42 and Rac1 gradients at cellular front during cell migration. We show that Cdc42 and Rac1 gradients are formed thanks to a combination of distributed GEFs and GAPs and not directly by diffusion or advection from a localized source. A minimal mathematical model suggests the following mechanisms: i) Cdc42 and Rac1 GEFs are exponentially distributed with a decay length of ~10μm; ii) the overall GAP activity for Cdc42 is uniform, such that the amount of active Cdc42 simply follows its GEFs distribution; iii) In addition to a uniform GAP activity, the Rac1 gradient is inhibited at the front by the β2-chimaerin GAP such that the Rac1 gradient extent (defined as the length scale at which Rac1 decreases by half) is increased. We show that the localized activity of β2-chimaerin depends on a negative feedback from both Cdc42 and Rac1, and that the actin retrograde flow is required for β2-chimaerin enrichment. Finally, we show that the resulting spatial properties of Cdc42 and Rac1 gradients govern the directionality and the speed of cell movement, respectively.

## Results

### Cdc42 and Rac1 gradients show two distinct shapes at the front of migrating cells

We investigated the spatial activity gradients of Cdc42 and Rac1 Rho GTPases at the basal plasma membrane by imaging FRET biosensors^40^. HeLa cells stably expressing FRET reporters were left to migrate randomly on glass coverslips, and were imaged using TIRF microscopy. The FRET ratio was calculated as a proxy for GTPase activity. Front-to-back gradients of either Cdc42 or Rac1 activity were measured from the cell protruding edge to the nucleus (**Fig 1a**). As previously reported in neutrophils^9^, we observed gradients that differed both in shape and in spatial extent. Cdc42 gradient was steep and monotonous, peaking at the protruding edge, and presenting an exponentially decaying profile of characteristic length d = 8.3 μm ± 0.6 μm (SEM, n=19). In contrast, Rac1 gradient peaked at a distance d = 5.8 ± 0.5 μm from the cell edge, and then decayed with a characteristic length d = 9.6 μm ± 0.7 μm (characteristic length of the exponentially decaying part, n=31). We defined the extent of the gradient by the distance between the tip of the cell and the point where the signal reaches half-amplitude. The extent for Rac1 was d = 14.6 ± 0.7 μm, compared to d = 8.9 ± 0.6 μm for Cdc42 (**Fig 1b**). Interestingly, these observations match those reported for gradients in other cell lines^9,41^. We thus questioned what could be the mechanisms underlying these gradients and accounting for their distinct shapes.

**Figure 1:**
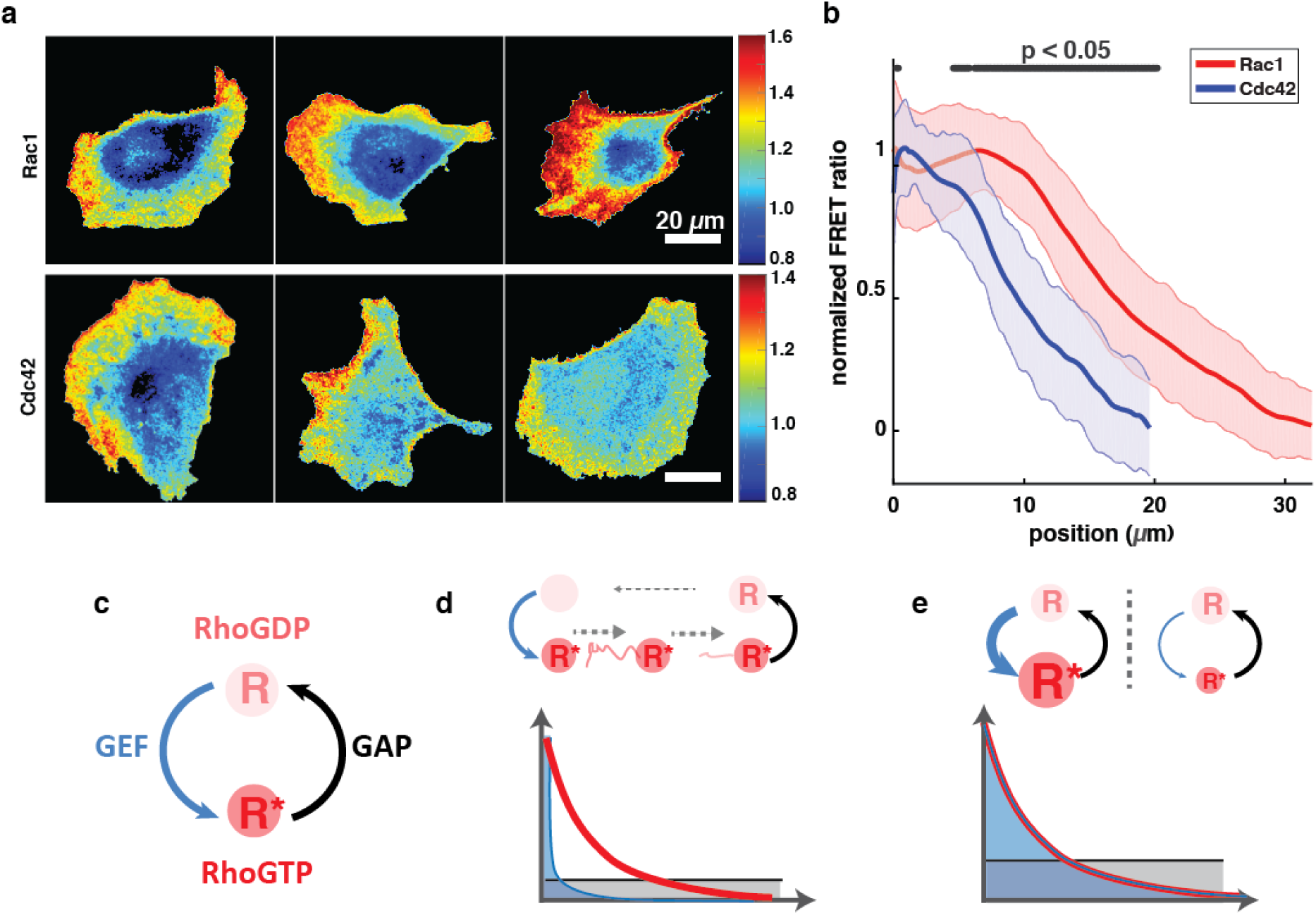
Rac1 and Cdc42 activity gradients have different shapes. **(a)** FRET biosensors were used to monitor Rac1 (top) or Cdc42 (bottom) activity in freely migrating HeLa cells. GTPase activity is measured by the FRET ratio, and represented with a color scale. Several representative cells are shown. Scale bar: 20 μm. **(b)** Normalized FRET ratio of Rac1 (red) and Cdc42 (blue) are plotted as a function of the distance from the cell edge. The error bars indicate the standard deviation (s.d) of n=31 (Rac1) or n=19 (Cdc42) cells. Black segments at the top show positions at which the curves are statistically different (p<0.05, Wilcoxon’s rank sum test) **(c)** Rho GTPase futile cycle, where the protein switch between an inactive and active state thanks to activators (GEFs) and deactivators (GAPs). **(d,e)** Two limit cases can be envisaged to explain the formation of cellular-scale Rho GTPase gradients. **(d)** A sharply localized GEF (blue profile) acts as a punctual source of active Rho GTPases (red) that are further transported by diffusion or advection (dashed grey arrows) until they reverse to the inactive state thanks to a GAP (black). **(e)** A cellular-scale distributed GEF activates locally the Rho GTPase such that both have the same profile.

Two generic classes of model can account for the patterning of spatially graded distributions^34^. The first class relies on transport mechanisms (diffusion, advection) to establish gradients from a localized source (**Fig 1c,d**). A canonical example is the synthesis-diffusion-degradation model (SDD), which has been heavily discussed in the context of the Bicoid morphogen gradient^42^. The second class of models assumes a graded distribution of activators and deactivators (**Fig 1c,e**). In this context, the local concentration is set by the local balance between activation and deactivation. This second class of model has also been proposed to explain the establishment of morphogen gradients, e.g. for the formation of the bone morphogenetic protein (BMP) gradient that patterns the dorso-ventral axis of the early *Xenopus* embryo^43,44^.

### Cdc42 and Rac1 gradients are shaped by spatially distributed GEFs and GAPs but not by diffusion

In order to distinguish between these two classes of model, we opted for an input-output relationship approach. We used optogenetics^45,46^ to impose activation gradients of either Intersectin-1 (ITSN) or T-Cell Lymphoma Invasion And Metastasis 1 (TIAM1), two GEFs specifically activating Cdc42 or Rac1, respectively. We used fusions of CRY2 with the DH-PH catalytic domain of ITSN or TIAM to activate specifically Cdc42 or Rac1^46^ (**Fig 2c**). A home-made illumination setup using a DMD (Digital Mirror Device^47^) allowed us to shine spatial gradients of light with an 8-bit gray level resolution. Cells were confined on round micropatterns to prevent cell shape polarity^48^ and gradients of light with slopes ranging from 1x to 4x were applied (**Fig 2a**). As we could predict in a previous work^49^, recruitment of the cytoplasmic optogenetic partner CRY2 to the basal plasma membrane followed the stimulation signal with the addition of an exponential decaying tail of 5 μm characteristic length due to the diffusion of CIBN-CRY2 dimers at the membrane (**Fig 2b**). This allowed us to tune precisely the spatial distribution of desired GEFs and test the relationship between the activation input and the output in terms of GTPase activity. If any transport mechanism (model 1) was taking place, we would expect a difference in the spatial distribution of the output compared to the input. For example, diffusion would smoothen the input, giving rise to a more extended distribution of the output by the addition of a length scale 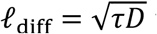 where τ is the lifetime of the Rho GTPase in its active GTP-bound state and *D* is its lateral diffusion coefficient. Contrarily, model 2 predicts that the spatial distribution of the output would mirror the distribution of the input, given that deactivators are uniform. Indeed, we reasoned that the optogenetic activation would dominate the other sources of activation such that the input-ouput relationship would reveal the distribution of the deactivators.

**Figure 2:**
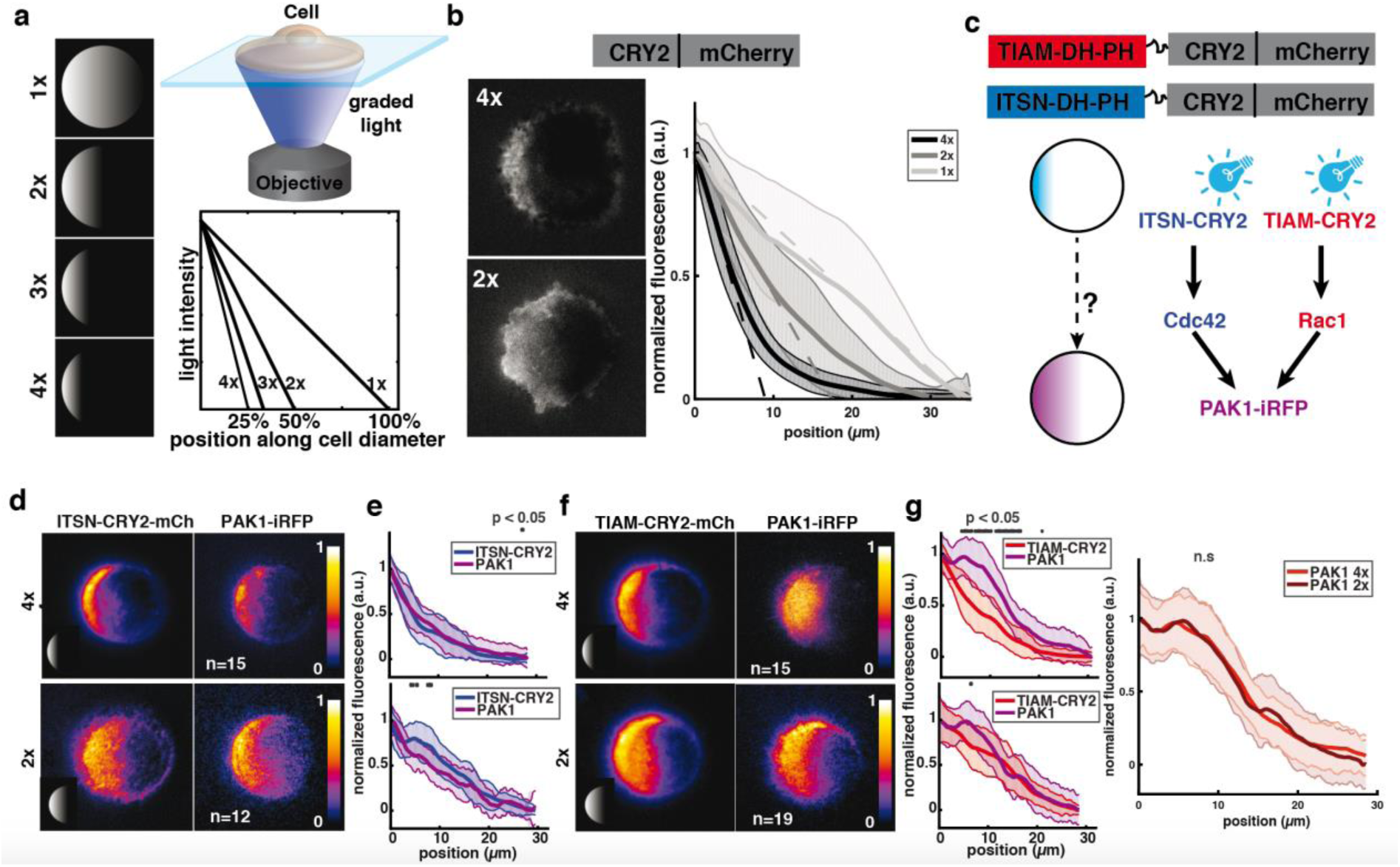
Cdc42 and Rac1 have different responses to GEF activation. **(a)** We imposed GEF activity gradients of different slopes using optogenetics. Patterned illumination with grayscale levels (left) was shone onto the samples, imposing linear light gradients of the same amplitude but different spatial extents on cells attached on round micro-patterns of 35 μm in diameter (top right). The reference gradient called “1x “spans over the whole diameter of the cell. The gradient “2x” spans over a half cell diameter, and therefore has a slope twice as sharp as for the gradient “1x” (bottom right). **(b)** Membrane recruitment of the optogenetic partner CRY2-mCherry to the basal side of cells on round micro-patterns following 30 minutes of illumination with 4x (top left) and 2x (bottom left) gradients. mCherry fluorescence (solid lines) was measured along the cell diameter following illumination with 1x (light grey), 2x (medium grey) and 4x (black) gradients (dashed lines). Error bars indicate the s.d. of n=15 (1x), n=24 (2x) and n=15 (4x) cells. **(c)** We activated GEFs of Cdc42 (ITSN) or Rac1 (TIAM) with light gradients and measured the fluorescence pattern of PAK1-iRFP. Schemes at the top represent the fusion proteins used. **(d)** PAK1-iRFP recruitment following 4x (top) or 2x (bottom) activation gradients of ITSN. Fluorescence was recorded using TIRFM in HeLa cells on round micro-patterns, and initial fluorescence was subtracted for normalization. Micrographs represent the averaged fluorescence of n=15 (4x, top) or n=12 (2x, bottom) cells. Insets show the illumination patterns (not to scale). **(e)** Normalized fluorescence of ITSN-CRY2-mCherry (blue) and PAK1-iRFP (purple) was measured along the cell diameter and averaged (solid lines). Error bars: s.d. Gray lines at the top show positions at which the curves are statistically different (p<0.05, Wilcoxon’s rank sum test). **(f)** PAK1-iRFP recruitment following 4x (top, n=15) or 2x (bottom, n=19) activation gradients of TIAM. **(g)** Normalized fluorescence of TIAM-CRY2-mCherry (red) and PAK1-iRFP (purple) along the cell diameter of n=15 (4x, top) or n=19 (2x, bottom) cells, Error bars: s.d. Gray: Wilcoxon rank sum test (p<0.05). Single cell data for **(e)** and **(g)** are presented in **supplementary figure 3**.

We thus sought to determine whether the GTPase activity of Cdc42 and Rac1 followed the activation pattern of their respective GEFs. Due to the difficulty to combine optogenetics and FRET imaging because of the overlapping wavelengths of imaging and activation, we used downstream effectors as reporters of GTPase activity. The protein PAK1 is activated downstream of both Cdc42 and Rac1. We thus monitored the basal membrane recruitment of a PAK1-iRFP fluorescent reporter following gradient activation of each one of the GTPases (**Fig 2c**). As controls, we verified that the observed recruitment of PAK1-iRFP was not due to fluorescence bleed-through or non-specific activity of CRY2-mCherry (**Supplementary Figure 1a**), nor to volume effects or cell deformation (**Supplementary Figure 1b**). Importantly, we also verified that the GEF DHPH domains used in our optogenetic approach were truly specific and did not observe any cross-activation (**Supplementary Figure 2**). PAK1-iRFP recruitment patterns followed the activation gradients of ITSN-CRY2 remarkably well, independently of their spatial extents (**Fig 2d,e, Supplementary Video 1**). We could not detect any significant difference between the activating ITSN gradient and the PAK1 response, up to the resolution of our measurement estimated as ~2μm (two standard deviations of the spatial extent). This result suggests that GEF activity levels are sufficient to shape Cdc42 activity patterns without the requirement of other mechanisms such as transport. More precisely, our result shows that the contribution of diffusion is bounded, ℓ_diff_ < 2μm, which sets an upper limit for the lifetime of Cdc42 in the GTP bound state to 2s assuming a usual lateral diffusion coefficient of 0.5μm^2^/s. Conversely, PAK1-iRFP spatial recruitment was independent of the shape of the activating TIAM-CRY2 gradient. It did not follow the sharpest activation gradient (4x), and the peak at 6 μm from the protrusion edge was present from the beginning of the stimulation (**Supplementary Figure 4, Supplementary Video 2**) despite its absence from the gradients of TIAM-CRY2 (**Fig 2f,g**). Interestingly, the PAK1-iRFP gradient obtained with our synthetic approach matched the Rac1 gradient observed in native cells (**figure 1b**). Thus, a more complex scenario is required in the case of Rac1 than for Cdc42. We sought to discriminate between two possibilities explaining how the Rac1 gradient is being shaped: whether shaping involves transport or non-uniformly distributed deactivators.

### A crosstalk between Cdc42 and Rac1 through GEFs and GAPs contributes to Rac1 gradient shaping

We investigated the molecular mechanisms shaping the gradient of Rac1. A complex crosstalk between the Cdc42 and Rac1 pathways has been shown previously^12,28^. We questioned whether such network could explain the complex pattern of Rac1 activity we observed. We used the Abi1-iRFP fusion protein as a reporter of Rac1 activity. Abi1 is part of the WAVE complex that has been shown to be activated specifically by Rac1 but not Cdc42^50^ (**Fig 3a**). We monitored Abi1-iRFP recruitment following optogenetic activation of either TIAM or ITSN. We observed that Abi1 is activated at the cell edge following TIAM but also ITSN activation (**Fig 3b**), suggesting that Cdc42 directly or indirectly activates Rac1. Interestingly, in both cases the induced Abi1 recruitment was more restricted to the cell border than the activating gradients (**Supplementary Figure 5**). This observation is in accordance with the known distribution of the WAVE complex at the tip of the lamellipodia^51^, suggesting a compartmentalization independent of the immediate Rho GTPase activation. Yet, in addition to the positive crosstalk, we also observed a cross inactivation of Rac1 by Cdc42. When we inhibited Cdc42 by siRNA (**Supplementary Figure 6a**), we observed an increase of Rac1 activity at the cell front as measured by the non-normalized FRET profile (**Fig 3c**). Strikingly, the bump of Rac1 activity 6 μm from the cell edge was abolished in this condition. Since the overall effect of Cdc42 depletion is to increase Rac1 activity, we reasoned that the dominant role of Cdc42 on Rac1 is to specifically activate a GAP inhibiting Rac1.

**Figure 3:**
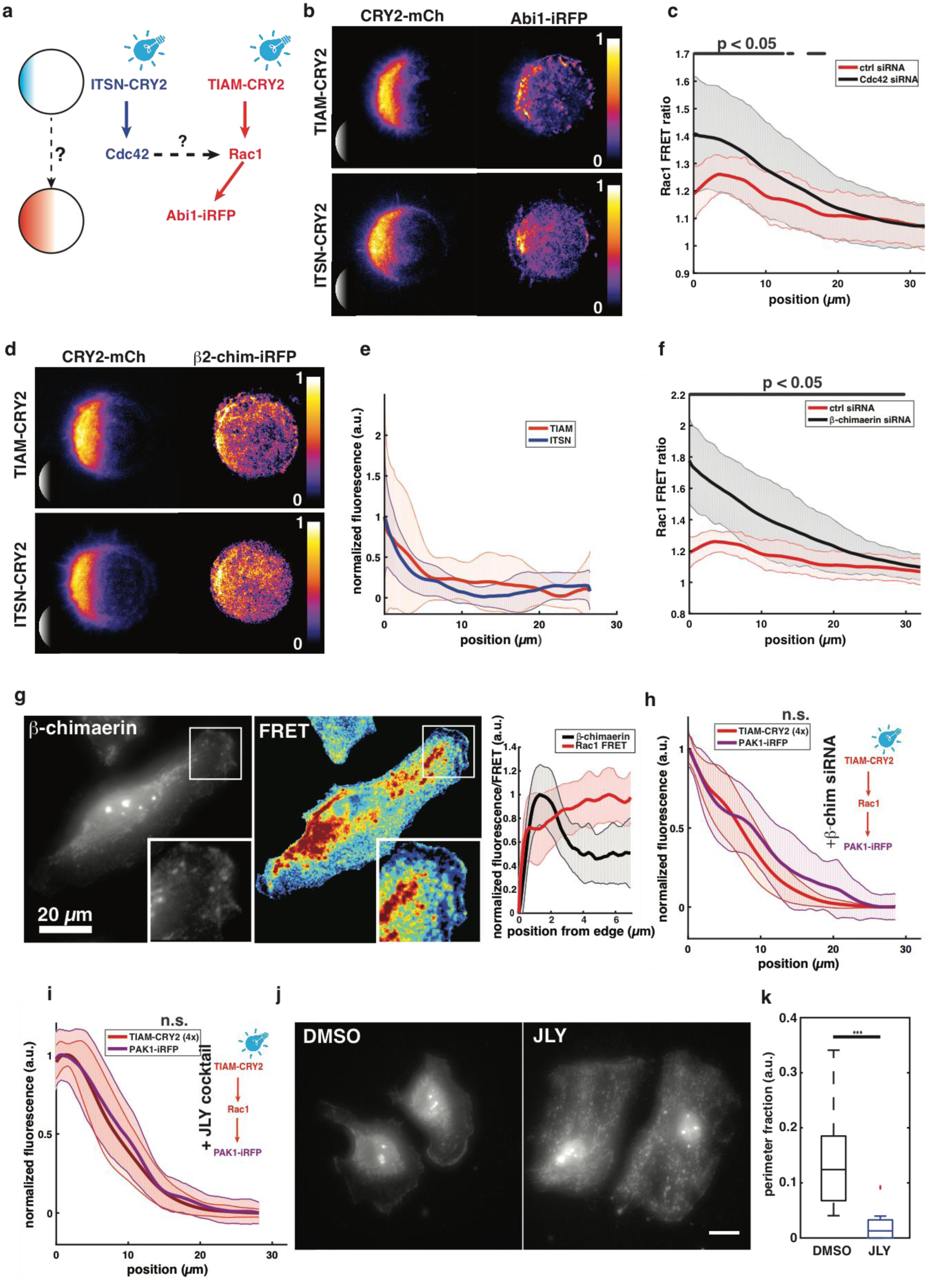
a crosstalk between Cdc42 and Rac1 involves β2-chimaerin to shape the activity gradient of Rac1. **(a)** We activated GEFs of Cdc42 (ITSN) or Rac1 (TIAM) with light gradients and measured the fluorescence pattern of Abi1-iRFP. **(b)** Averaged Abi1-iRFP recruitment (right column) following 4x activation gradients (left column) of TIAM (n=10, top) or ITSN (n=11, bottom) visualized using TIRFM on round micro-patterns. The averaging procedure is explained in the methods section. Insets show the illumination patterns (not to scale). **(c)** Non-normalized Rac1 FRET ratio profiles along cell diameters of cells treated with control siRNA (n=38, red) or Cdc42- directed siRNA (n=37, black). Error bars: s.d. Gray lines at the top show positions at which the curves are statistically different (p<0.05, Wilcoxon’s rank sum test). **(d,e)** β2-chimaerin-iRFP (right) recruitment following 4x activation gradients (left) of TIAM (n=11, top) or ITSN (n=12, bottom), imaged in TIRFM on round micro-patterns. **(d)** Micrographs represent the averaged fluorescence (see methods). **(e)** Normalized fluorescence of β2-chimaerin-iRFP was measured along the cell diameter following the activation of ITSN-CRY2-mCherry (blue, n=12) or TIAM-CRY2-mCherry (red, n=11). Error bars: s.d. **(f)** Non-normalized Rac1 FRET ratio profiles along cell diameters of cells treated with control siRNA (n=38, red) or β2-chimaerin -directed siRNA (n=25, black). Error bars: s.d. Gray: Wilcoxon rank sum test (p<0.05). **(g)** β2-chimaerin staining (left, Gamma correction was applied to images in order to visualize the full dynamics) compared to normalized Rac1 FRET (middle) in the same cells. Insets show zoomed regions of the cell edge. Levels of β2-chimaerin (black) and Rac1 activity (red) are anti-correlated at the cell front (right panel, n=8, Error bars: s.d). **(h)** PAK1-iRFP (purple) recruitment following 4x activation gradients of TIAM (red) after treatment with β2-chimaerin directed siRNA (n=14). Curves were found not significantly different on their whole length (Wilcoxon, p> 0.05). **(i)** PAK1-iRFP (purple) recruitment following 4x activation gradients of TIAM (red) after treatment with JLY cocktail (n=23).n.s.: non-significant. **(j)** β2-chimaerin staining after DMSO (left) or JLY cocktail (right) treatment. **(k)** Fraction of cell perimeter showing β2-chimaerin signal at the cell edge larger than in the cytosol (DMSO: n=11, JLY: n=10). Fluorescence at the cell edge was measured along a 1 micrometer-thick line obtained from the thresholding-based segmentation of the cell shape. The signal in the cytosol was evaluated from a 1 micrometer-thick line outlining that cell edge on its cytosolic side. Boxplots represent the median, interquartile (box), 1.5 IQR (whiskers) and outliers (red crosses). Statistical significance was evaluated using Wilcoxon’s rank sum test. ***: p ≤ 0.001

β2-Chimaerin is a GAP of Rac1 that was shown to be activated at the protrusion edge downstream of chemotactic signals^52^. We monitored the recruitment of the β2-chimaerin-iRFP reporter following the optogenetic activation of TIAM or ITSN. We could observe that both pathways could recruit β2-chimaerin at the cell edge, in a very localized fashion similar to the WAVE recruitment (**Fig3 d,e**), suggesting that this GAP recruitment is conditioned by other signaling components belonging to the tip of the lamellipodia. Accordingly, inhibiting β2-chimaerin using siRNA (**Supplementary Figure 6b**) led to a strong increase of Rac1 activity especially at the cell front such that the bump was abolished (**Fig 3f**). This result suggests that β2-chimaerin might act downstream of Cdc42 and Rac1 to inhibit Rac1 locally at the cell front. Indeed, at the front of randomly migrating cell we observed an anti-correlation between β2- chimaerin and Rac1 activity measured by FRET (**Fig 3g**), where β2-chimaerin peaks at the cell edge and Rac1 activity increases when β2-chimaerin decays. We could verify that the observed localization of β2-chimaerin at the cell edge was not due to volume effects related to the local membrane ruffling activity (**Supplementary Figure 7**). We further confirmed the direct role of β2-chimaerin in shaping the Rac1 gradient by inducing the sharp Rac1 activation (4x) using optogenetics in β2-chimaerin depleted cells, which resulted in a PAK1 gradient that now matched the activating profile (**Fig 3h**).

Orthogonally to the previous experiments, we also tested the role of transport in shaping the Rac1 gradient. Given that we observed the same PAK1 spatial profile for two distinct TIAM-CRY2 activating gradients (**Fig 2g**), we excluded diffusion as we would have observed two smoothened output distributions if diffusion was acting on it. Conversely, advection due to the retrograde flow of actin in the lamellipodia can give rise to two similar outputs if the distribution of the flow velocities is ultimately limiting for the spatial expansion of the gradient. When cells were treated with the Jasplakinolide-LatrunculinB-Y27632 (JLY) drug cocktail that freezes actin dynamics^53^, we indeed observed a Rac1 activity gradient that matched the sharp (4x) TIAM-CRY2 input gradient (**Fig 3i**). Surprisingly, this result shows that the actin retrograde flow can also account for the bump observed in the endogenous Rac1 gradient besides our previously found role for β2-chimaerin. However, this effect could be indirect if the retrograde flow was acting not on Rac1 itself but on the machinery required for proper β2-chimaerin localized distribution. To test this hypothesis, we compared the distribution of β2-chimaerin in control and JLY-treated cells (**Fig 3j**). β2-chimaerin localization disappeared from the tip of migrating cells in JLY-treated cells, confirming the indirect role of actin dynamics. (**Fig 3k**).

### A minimal model of local reactions recapitulates Cdc42 and Rac1 gradient shaping

Given the numerous layers of interactions that we identified experimentally, we sought for a minimal model that would capture the main mechanisms giving rise to the cellular scale properties of the Cdc42 and Rac1 gradients. To this end, we built a one-dimensional model, where the *x*-axis spanned across the cell from *x*=0 to *x*=35μm. We assumed that the Rho GTPases were activated and deactivated with first order kinetics, and that levels of Rho GTPases equilibrated on a fast time scale. We assumed that the total amount of Rho GTPase *R*_tot_ was not limiting. Eventually, we excluded diffusion and advection, such that the model was purely local. Thus, the local concentration of active Rho GTPase *R*\*(*x*) at steady-state is of the form:

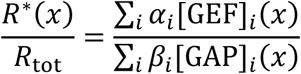

where [GEF]_*i*_(*x*) and [GAP]_*i*_(*x*) are the concentration profiles of GEFs and GAPs, and *α_i_* and *β_i_* the effective activation and deactivations rates (which can be a function of the concentration of the RhoGTPases themselves in the case of crosstalks). The sums over *i* go over all possible sources of activation and deactivation (see **Fig 4a** for all terms based on our experimental findings). We built a numerical tool to screen for the different roles of all previously identified parameters such that we could extract a minimal model for the formation of Cdc42 and Rac1 gradients (**Fig 4b**). For Cdc42, the shape of the gradient can be simply explained by an exponentially distributed GEF and uniform GAP (**Fig 4c**:

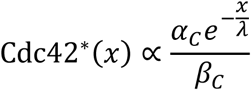

where λ is the decay length measured for Cdc42 itself (about 10μm). Note that in the case of optogenetic activation, the optogenetic term 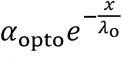 most probably dominates the endogenous GEF activity (*α*_opto_ ≫ *α_C_*) such that the induced gradient follows the activating one. For Rac1, our model contains: i) an exponentially distributed GEF of 10μm decay length and uniform GAP, similarly to Cdc42. The values of *α_R_* and *β_R_* for Rac1 relative to the ones of Cdc42 were chosen to match the relative FRET levels of Rac1 and Cdc42 (**Supplementary Figure 4**), were Rac1 activity is about 3 times larger than Cdc42 while presenting the same characteristic length in its exponentially decaying tail (*α_R_* = 2*α_C_* and *β_R_* = ½ *β_C_*). ii) In addition to the uniform GAP, a second one (namely *β*2-chimaerin) was assumed to be exponentially distributed with its own characteristic length γ = 5μm. With these ingredients, Rac1 takes the following form:

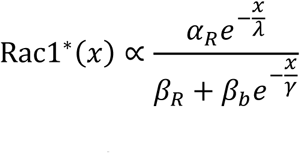

**Figure 4:**
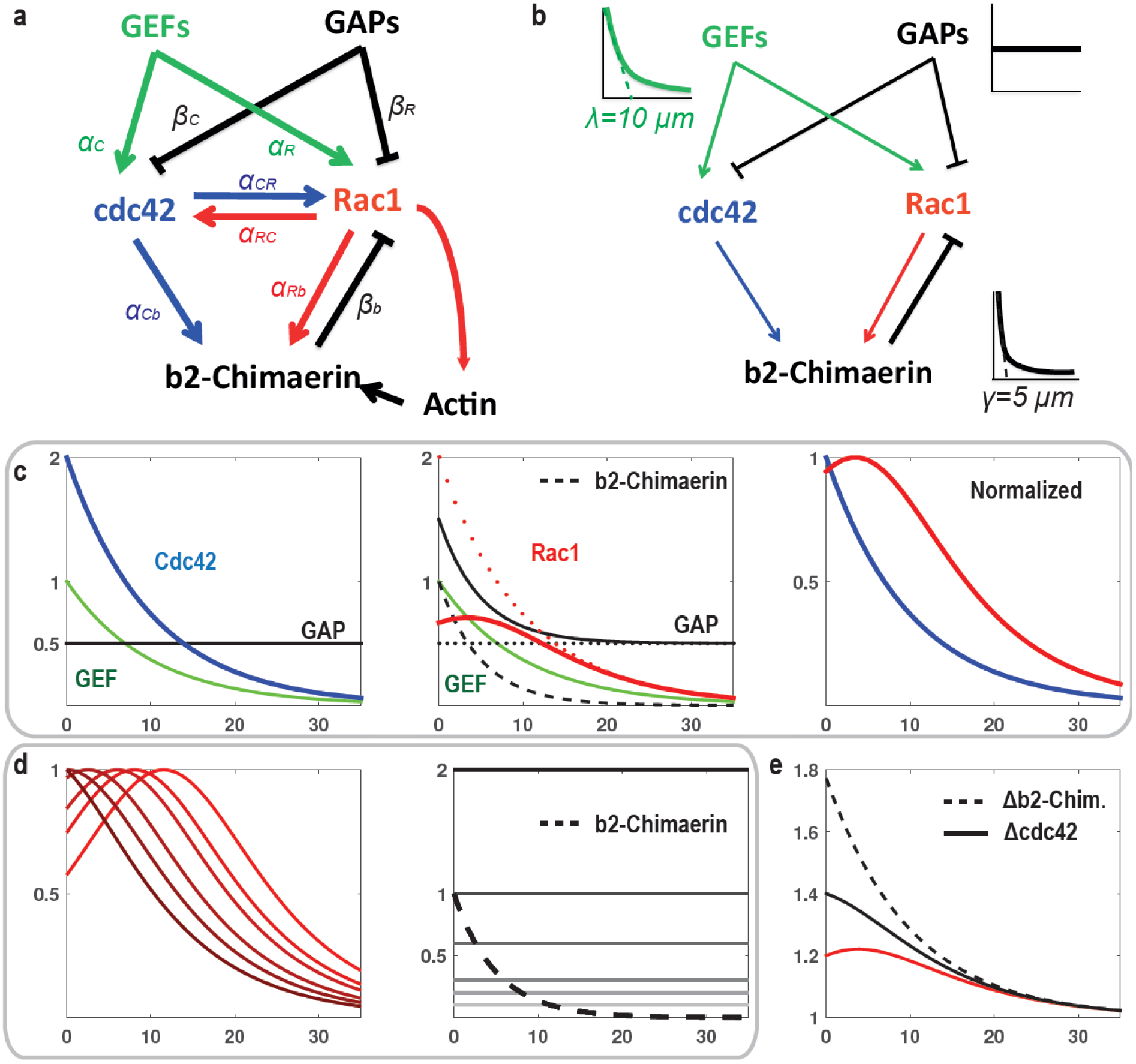
a minimal model for Cdc42 and Rac1 gradient formation. **(a)** Full model based on our experimental findings. Activation rates are denoted by α and deactivation rates by β. **(b)** A minimal model that recapitulates the formation of Cdc42 and Rac1 gradients. The thickness of the arrows is representative of the strength of the links. **(c)** Gradient shaping of Cdc42 and Rac1. Since the absolute amplitude of Rho GTPases are unknown, we assigned the following arbitrary values to the rates: α_C_ = α_R_ = 1; β_C_ = β_R_. = 0.5; and β_b_ Left: Cdc42 (blue) is set by an exponentially distributed GEF (green) with a characteristic length λ=10μm and uniform GAP (black). Middle: Rac1 (red) requires an additional GAP, β2-Chimaerin (dashed black, characteristic length γ=5μm), that is localized more sharply at the cell edge than Rac1 GEF (green). The overall GAP activity (plain black) is the sum of β2-Chimaerin and a uniform GAP (dashed black). As a result, the putative Rac1 gradient without β2-Chimaerin (dashed red) is chopped off at the cell edge resulting in a bell-shaped gradient (plain red). Right: once normalized to 1, Cdc42 and Rac1 gradients present distributions that are similar to the ones measured in cells. **(d)** Effect of the relative ratio r = β_R_/β_b_ between uniformly distributed GAPs and the localized gradient of β2-Chimaerin on the position of the Rac1 bump. Left: Rac1 gradients obtained with decreasing values of r (r = 2, 1, 0.6, 0.3, 0.2, 0.1 respectively from dark to light red). Right: exponentially distributed β2- Chimaerin (dashed line) and uniformly distributed GAPs =(β_R_ 2, 1, 0.6, 0.3, 0.2, 0.1 from dark to light gray, solid lines) corresponding to the values used for the left plot. Right: sum of the GAPs shown in the middle plot. **(e)** Effects of Cdc42 or β2-Chimaerin inhibition in silico on the Rac1 gradient (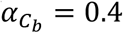 and 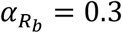) The profiles are normalized (by the same factor) to match the FRET signal values measured experimentally (**figure 2c,d**).

Where *β_b_* is the effective rate constant for β2-chimaerin GAP activity on Rac1. This expression for Rac1 is sufficient to explain the bump (**Fig 4c**), the position of which is determined by the ratio *r* = *β_R_*/*β_b_* between the strength of the uniform GAP over the strength of the localized β2-chimaerin, and by the characteristic lengths of the decaying profiles:

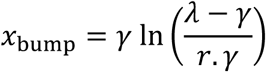

From this relationship, we can see that a bump will be present if *r* < (*λ* – *γ*) /*γ* which reduces to *r* < 1 using the experimental numbers for the decay lengths, or equivalently *β_R_* < *β_b_* This means that the strength of the uniform GAP has to be less than the strength of β2- chimaerin to observe a Rac1 bump. The evolution of the bump position as a function of *r* is presented in **Fig 4d**. From the bump position observed in our experiment, we could predict that 2-chimaerin dominates the uniform GAPs by a factor of ~ 2. This minimal model for Rac1 can be refined to account for the respective roles of Cdc42 and Rac1 in mediating *β*2- chimaerin activity at the tip (**Fig 4e**). Assuming that is a linear function of Cdc42 and Rac1 concentrations: *β_b_* = *β_cb_* Cdc42(*x*) + *β_Rb_* Rac1(*x*) the model shows that Rac1 self-inhibition is required to account for the observed differences in the Rac1 gradient between cells depleted for Cdc42 and cells depleted for β2-chimaerin (*β_Cb_* = 0.7 = 2*β_Rb_*). Altogether, our (minimal) modeling approach suggests a simple mechanism of distributed activators and deactivators that shape Cdc42 and Rac1 gradients such that their spatial extents are ultimately different. Compared to Cdc42, Rac1 gradient is extended thanks to a sharply distributed deactivator. We thus anticipated that the spatial extent of these Rho GTPases would play a functional role.

### A controlled assay to monitor the dependence of cell migration on Rac1 and Cdc42 gradients

We questioned whether the different properties of Cdc42 and Rac1 gradients had an impact on migration properties. Using optogenetics, we thus tested how gradients of ITSN or TIAM with increasing slopes (**Fig 2a**) would affect cell polarization and migration. In order to control the experimental initial conditions, i.e. to prevent initial cell polarity prior to the optogenetic stimulation but still be able to monitor cell movement following it, we opted for a switchable micropatterning technique^54^. Cells were plated on round micropatterns, and would then keep an isometric shape until the surrounding repelling surface was rendered adhesive by coupling a fibronectin-mimicking chemical compound (BCN-RGD) that binds to the modified PLL-PEG repellent (APP). After addition of this reagent, cells were released from patterns and free to migrate on the coverslip (**Fig 5a-b**, top row). Optogenetic stimulation with gradients of light concomitantly with the release of adhesion allowed us to study cell migration with one changing parameter, namely the extent of blue light gradients (**Fig 5a-b**, bottom row). From n~20 cells per each condition, we quantified the cell edge morphodynamics (**see methods**) and averaged them for each activating gradient slope (**Fig 5c**). As expected, both Rac1 and Cdc42 biased the membrane protruding activity toward the direction of the gradient. Rac1 led to an immediate cell movement while Cdc42 led to slightly delayed cell movement (**Fig 5c**). We observed that cells shifted from an oriented spreading (when the back of the cell kept steady) to a directed migration (when the back of the cell moved together with the front) by increasing the gradient slope (**Supplementary Video 3, 4**). Yet, the center of mass of cells monotonously increases its movement toward the gradient as the gradient slope increased (**Fig 5d**) suggesting that the quantitative properties of the gradients have a differential role in migration.

**Figure 5:**
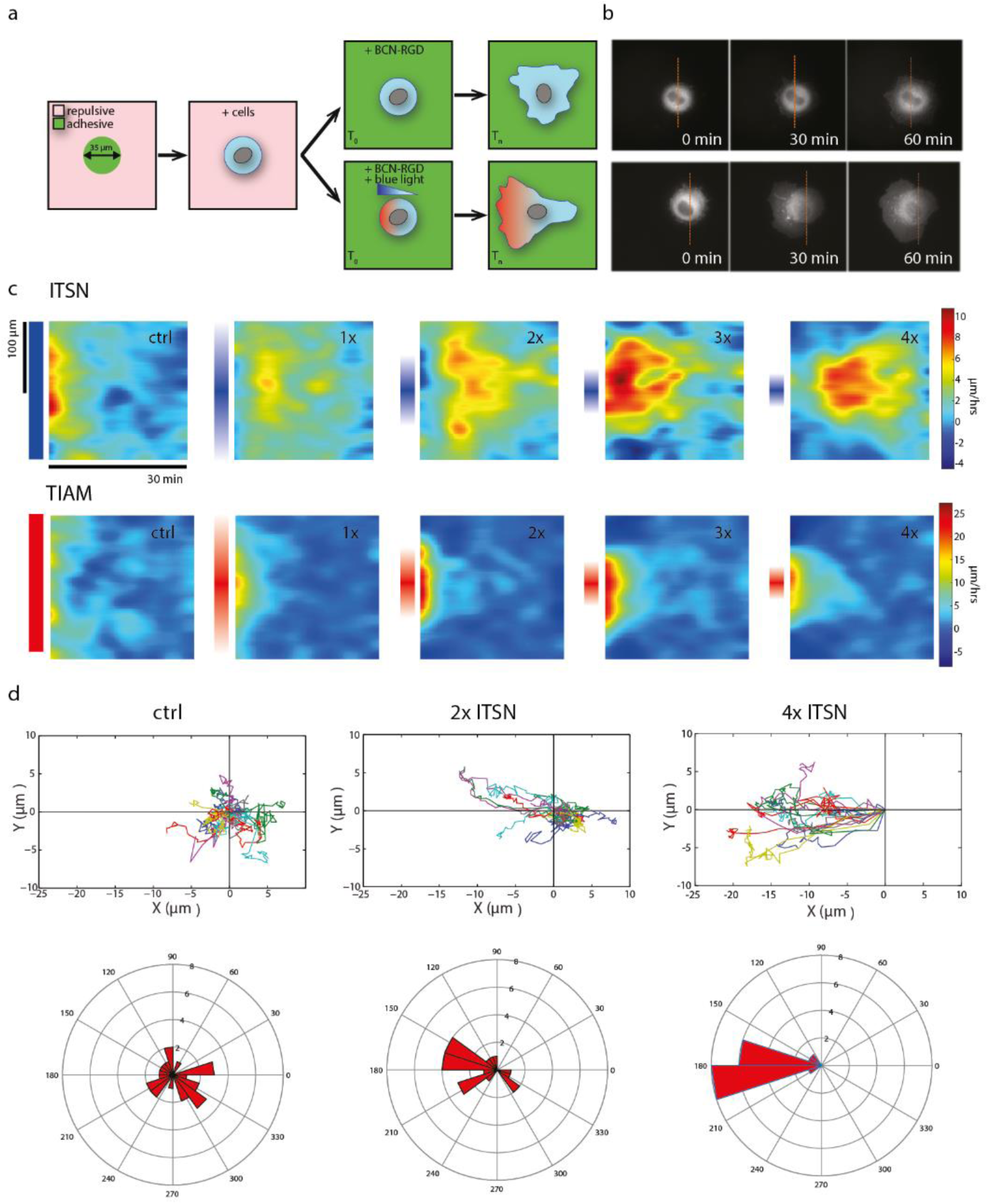
scheme of the quantitative migration assay. **(a)** Cells are seeded on 35μm round patterns. After complete adhesion, the adhesive reagent BCN-RGD is added and binds to the coverslip’s surface, allowing free 2D cell-migration (top). Directed migration can be triggered by optogenetic activation of GEFs through light gradients at the same time as cell adhesion is released (bottom). **(b)** Examples of cells expressing CIBN-GFP-CAAX and TIAM-CRY2-mCherry with (3x gradient, bottom) or without (top) photo-activation (visualized: TIAM-CRY2-mCherry). Time indicates the duration after addition of BCN-RGD and concomitant blue light illumination. The dashed orange line corresponds to the initial position of the cell center. **(c,d)** HeLa cells expressing CIBN-GFP-CAAX and ITSN-CRY2-mCherry or CIBN-GFP-CAAX and TIAM-CRY2-mCherry were illuminated with various gradients of light as the adhesive patterns were released. **(c)** Average morphodynamic maps for each condition (ITSN: top, TIAM: bottom). The vertical axis corresponds to the coordinate along the cell contour (centered on the direction of the light gradient) and the horizontal axis corresponds to time. The local velocity of the edge of the cell membrane are color coded accordingly to the bar on the right side. Gradient extents are schemed on the left side of each map. ITSN: n=25 (control with uniform illumination), n=16 (1x gradient), n=20 (2x), n=19 (3x) or n=16 (4x). TIAM: n=19 (ctrl), n=18 (1x), n=18 (2x), n=17 (3x), n=18 (4x). **(d)** We tracked the position of the centroid of individual cells. Top: Trajectories of cells stimulated with various gradients of ITSN. Bottom: The angles between the displacement vector (initial to final centroid position) and the stimulation axis for each cell are represented in polar coordinates.

### Cdc42 provides directionality while Rac1 provides speed

In order to assess the quantitative effect of gradient on motility, we focused on two coarse-grained parameters: the maximal instantaneous velocity and the precision of the migration orientation. Sharper gradients of either ITSN or TIAM both increased cellular speed. However, activating Rac1 through TIAM had a stronger effect on speed than activating Cdc42 through ITSN (**Fig 6a**), consistent with the known effect of Rac1 as a critical factor for lamellipodium formation^15^ (**Supplementary Figure 8**). In fact, even shallow gradients of TIAM induced an enhanced migration speed. In comparison, only sharp gradients of ITSN (3x and 4x) induced an increased cellular speed, but even in these conditions speed was lower than for equivalent TIAM gradients. Conversely, ITSN gradients had a stronger effect on orientation precision. While 1x to 4x TIAM gradients had a similar effect on orientation, increasingly sharp gradients of ITSN induced an increasing precision of migration (**Fig 6b**). Indeed, the 4x ITSN gradient induced the most oriented response (with a remarkable angular precision), consistent with the known role of Cdc42 as a regulator of directed migration^55,56^, even though this role seems to be cell dependent^57^. We show here that directed migration is better achieved with sharp Cdc42 gradients similar to the ones measured endogenously in cells (Cdc42 gradient extent measured in migrating cells d = 8.9 ± 0.6 μm, **Fig 1**, 4x Cdc42 gradient extent imposed and measured through PAK-iRFP d = 6.1 ± 0.9 μm, **Fig 2**). Thus, in our experimental model, Cdc42 provides directionality while Rac1 provides speed of movement. These functions appear to be specific of each GTPase, since inhibition of Rac1 abolishes cell speed but not orientation (**Fig 6c,d**). Consequently, crossed activities (speed induction by Cdc42, orientation by Rac1) seem to be due to crosstalks between these RhoGTPases. Along this line, a possible functional role for β2-Chimaerin is to spatially segregate Rac1 and Cdc42 activities to avoid competition between their functional roles. Indeed, as seen in **supplementary figure 9**, β2-Chimaerin suppression has no effect on cell speed but leads to a significant reduction in angular precision. This suggests that β2-Chimaerin limits Rac1 protrusive activity at the very cell front to allow Cdc42 activity to steer cell migration.

**Figure 6:**
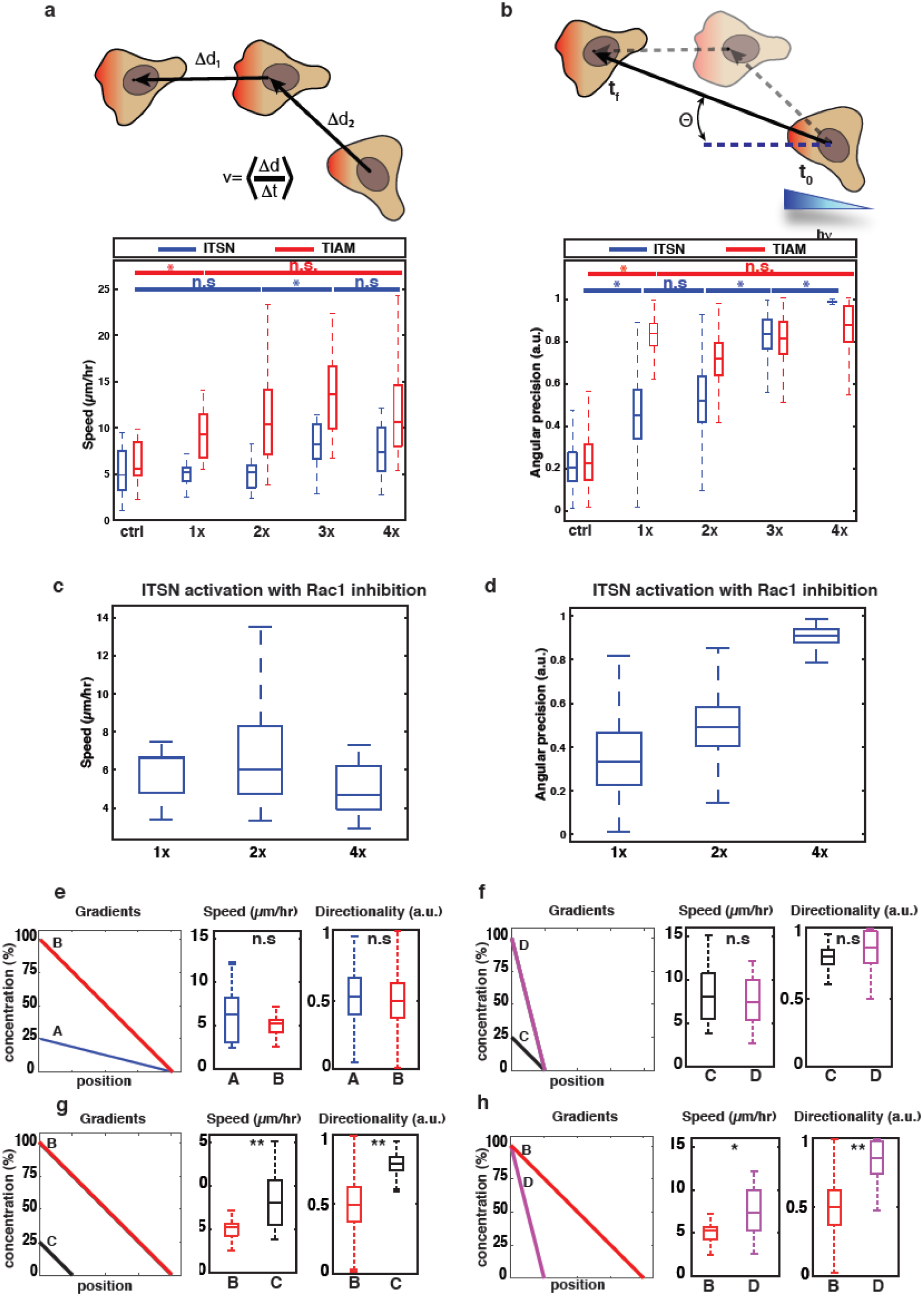
Cdc42 and Rac1 drive different cellular responses. **(a,b)** Trajectories of cells stimulated as represented in **Fig 4** were analyzed quantitatively. **(a)** Box plots showing the instantaneous speed of cells expressing CIBN-GFP-CAAX together with ITSN-CRY2-mCherry (blue) or TIAM-CRY2-mCherry (red) stimulated with various gradients of light (ITSN: n=25 (ctrl), n=16 (1x), n=20 (2x), n=19 (3x), n=16 (4x), TIAM: n=19 (ctrl), n=18 (1x), n=18 (2x), n=17 (3x), n=18 (4x)). For each cell, we averaged the instantaneous speed measured between consecutive time frames (top scheme). Boxplots represent the median, interquartile (box), 1.5 IQR (whiskers). **(b)** Angular precision of cell displacement. Angles measured as in (a) were bootstrapped, and angular precision was calculated with the formula 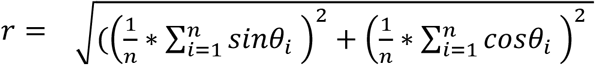. Boxplots represent the median, interquartile (box), 1.5 IQR (whiskers). Statistical significance between consecutive conditions was evaluated using Wilcoxon’s rank sum test. *: p ≤ 0.05. n.s. : non significant (p > 0.05). **(c,d)** HeLa cells expressing CIBN-GFP-CAAX and ITSN-CRY2-mCherry were stimulated with various light gradients after Rac1 inhibition with 100 μM NSC 23766, and cell motility was measured. **(c)** Box plots showing the instantaneous velocity in response to various ITSN gradients. n= 8 (1x gradient), n= 16 (2x gradient) or n= 17 (4x gradient). Boxplots represent the median, interquartile (box), 1.5 IQR (whiskers). **(d)** Angular precision of cell displacement. **(e-h)** Effects of slope vs. spatial extent of Cdc42 gradients on cell migration. HeLa cells expressing CIBN-GFP-CAAX and ITSN-CRY2-mCherry were stimulated with light gradients of varying slope, amplitude and spatial extent (y-axis represents relative concentrations tuned by modifying the light intensity). Cell velocity (speed, as in (a)) and directionality (angular precision as in (b)) were compared. **(e, f)** A (blue) and B (red) gradients have the same gradient extent but different slopes and amplitudes, as do C (black) and D (magenta). n = 13 (A), n = 16 (B), n = 22 (C), n = 16 (D). **(g, h)** B and C gradients have the same slope but different gradient extents and amplitudes, while B and D gradients have the same amplitude but different gradient extents and slopes. *: p ≤ 0.05,** : p ≤ 0.01 (Wilcoxon rank sum test).

### The spatial extent but not the amplitude or slope of the Cdc42 gradient matters for directionality

Since we showed that the shape of Rho GTPase activation gradients directly influence the outcome of cell migration, we thus questioned whether cells are actually sensitive to the slope or to the spatial extent of Rho GTPase activation gradients. In fact, in the previous experiments, both parameters varied concomitantly. It is known that cells can sense and process various extra- and intracellular signaling gradients that can hence influence cell polarity and migration^58–60^. However, it is not known to which quantitative properties of Rho GTPases intracellular signaling gradients cells are sensitive. Using the experimental setup detailed above, we could independently test the effect of gradient slope or spatial extent. When we applied gradients of ITSN activation with different slopes but the same spatial extent, we could not detect any difference in cell motility (**Fig 6e,f**). This also confirms that the amplitude of the imposed gradient itself does not affect the cellular response. Instead, when we imposed gradients of similar slope or amplitude but different extents, we could observe that cells stimulated with the shorter gradient of ITSN activation migrated with higher velocity and better orientation (**Fig 6g,h**). These results indicate that the spatial extent is the critical parameter of Rho GTPase gradients read by cells.

## Discussion

In this work, we reveal important aspects of the interaction network between Cdc42, Rac1 and their regulators, which led us to propose a network topology that is sufficient to recapitulate the spatial properties of these Rho GTPases activity gradients at the front of migrating cells. Using optogenetics allowed us to quantitatively tweak the input (spatial patterns of specific GEF activity for either Cdc42 or Rac1) and quantify the activity output, as measured by the recruitment of downstream effectors. By combining experimental and model-based approaches, we could spatially map the circuitry of the network and explain how intracellular patterns of Cdc42 and Rac1 are formed.

Cdc42 activity gradients can be simply explained by the combination of a localized GEF and a uniform GAP. Conversely, Rac1 patterning requires a more complex circuitry. Rac1 gradient has a complex shape, peaking at approximately 6 μm from the cell edge, and then decaying exponentially, whereas Cdc42 decays exponentially from the cell front. The spatial properties of these gradients are similar to what has been observed before in other cell types^9,41^. We experimentally find that two mechanisms could explain these observations. Combining one exponentially decaying GEF with either a GAP with a shorter exponential decay (like β2- chimaerin) or advection from the cell front is sufficient to recapitulate the observed Rac1 gradient. Yet, we show that the effect of advection does not act directly on Rac1 itself but is required for the front-most localization of β2-chimaerin. Indeed, in β2-chimaerin-depleted cells, the Rac1 activity follows its GEF distribution (**Fig 3h)**. Thus, actin treadmilling appears to be required for the localization of β2-chimaerin at the tip of the cell. It has been previously demonstrated that advection through the actin flow is coupled to cell polarity, by transporting various proteins away from the cell front^58^. In the present case, since β2-chimaerin is enriched at the cell front, either the actin flow act on an inhibitor of β2-chimaerin or there is another mechanism in action. One possibility is that β2-chimaerin localizes at the barbed end of actin filaments thanks to its interaction with the adaptor protein Nck1^52^. Nck1 is also localized at the tip of migrating cells by the Gab1-NWASP complex^61^. Since we additionally show that a feedback from Rac1 leads to β2-chimaerin enrichment, β2-chimaerin recruitment would depend on two concomitant signals: a Rac1 dependent signal likely going through Rac1-dependent PKC-DAG production^62^, and an actin polymerizing signal through the adaptor protein Nck1. This would also explain the crosstalk from Cdc42 to β2-chimaerin through N-WASP and an increase of Nck1-mediated β2-chimaerin recruitment.

Interestingly, Cdc42 and Rac1 gradients have similar exponential decays but different spatial extents due to the local inhibition of Rac1 activity at the cell front. This observation raises important questions about the way cells interpret signaling gradients, and about the parameters of these gradients actually affecting cellular response. These questions are also fundamental at the multicellular scale^63,64^. Using quantitative optogenetics, we could directly control the spatial extent, slope or amplitude of intracellular activity gradients. This strategy can be compared with the recent use of long-range light gradients and optogenetic activation of Cdc42 and Rac1 to guide cell migration^65^. Although our results suggest similar conclusions, our approach differs in its conception. In the work of Zimmerman and colleagues, optotaxis experiments rely on activation gradients that mimic the slopes of external chemo-attractant gradients, whereas we imposed intracellular activity gradients that mimic the ones measured endogenously^9^. We are convinced that it is more biologically relevant to consider the cell rather than the environment as the spatial referential for Rho GTPases gradients given that these gradients are set autonomously. Using our method, we showed that cell migration is not determined by the amplitude or slope of Rho GTPase gradients, but rather by their spatial extent, similarly to what was proposed in a recent work on ERK morphogen gradients in *Drosophila* embryos^64^. Thus, intracellular gradients may be processed in a different manner than the extra-cellular gradients, for which the relative slope matters thanks to mechanisms providing adaptation^66^. The spatial extent of Cdc42 needs to be small to ensure fine directionality in cell movement, in accordance with the previously shown role of Cdc42 as the primary conductor of chemotactic steering and cell polarity^9^. The spatial extent of Rac1 is larger, providing speed to the cell. Yet, since we could not effectively apply sharper Rac1 gradients without disrupting the network topology, we do not know if the spatial extent of Rac1 presents a functional optimum as for Cdc42.

In this study, we did not consider the temporal dynamics of Rho GTPase activities. While it is very likely that spatial and temporal dynamics are connected in freely migrating cells and while Rho GTPases patterns evolve on timescales of ~100s^19^, the response functions we measured under our steady optogenetic activations did not show evolving spatiotemporal patterns (**Supplementary Figure 4**). Thus, even if our synthetic approach does not recapitulate the full spatiotemporal complexity seen in native cells, we can consider our results as a signature of the signaling network capacity to respond to spatially modulated inputs. Given the similarity between the native and induced gradients of Rac1/Cdc42, we can be confident that the mechanisms we propose for gradient shaping are biologically significant, at least at the coarse-grained cellular scale.

Following a correlative approach, Yamao et al. recently studied in time and space the patterns of Rac1 and Cdc42 activities and their link with the membrane dynamics in randomly migrating cells^67^. They concluded that Cdc42 induces random cell migration and Rac1 is responsible for persistent movement. While it might sound different from our results, the discrepancies might be explained by the scales and parameters observed in each case. We measured local and instantaneous quantities (speed and directionality), and Yamao and colleagues measured integrated and macroscopic ones (persistence and randomness). We were not able to measure those integrated quantities, since our optogenetic activations were not following the cells as they moved out of the adhesive micro-patterns. However, these different scales can be reconciled. As we concluded that Rac1 provides cells with higher speed, it also means long-term movement is more persistent^58,68^. Similarly, since we showed that sharp Cdc42 gradients can fine-tune directionality, local and transient Cdc42 pulses (as measured at the cell edges in that paper) should steer cells randomly in complex trajectories. Moreover, our data fit with the tug-of-war model they propose: under such a model, the extended shape of Rac1 gradients should induce the formation of large protrusions, in line with the large lamellipodium usually attributed to Rac1 activation. Similarly, sharper, more local Cdc42 gradients should generate smaller protrusions in accordance with the Cdc42 activity usually associated with filopodia and small protrusions.

The minimal circuitry that we identified as sufficient to shape Cdc42 and Rac1 gradients raises new unanswered questions. In particular, it is unclear how gradients of GEFs and GAPs are shaped throughout the cell, beside the formation of the β2-chimaerin gradient we identified. Our results suggest a role for the cytoskeleton itself and its dynamics to enrich β2-chimaerin at the cell border. More generally, actin networks and actin-regulating complexes can act as scaffolding complexes in protrusive regions where they localize. For example, the WAVE Complex, a downstream effector of Rac1, recruits WRP, a GAP inhibiting Rac1^69^. Similarly, N-WASP, a downstream effector of Cdc42, associates with the GEF ITSN. More mechanisms are probably involved. In particular, membranes could play a direct role in the localization of these regulators of Rho GTPase activity. The local lipid composition, and in particular the concentration of PIP3, has been shown to control the activity of Rac1 and Cdc42^26,32,70^. In addition, membrane curvature-sensing BAR proteins localize at highly bent membranes, including cell edges. Several BAR proteins are known to bind Rho GTPAses or their regulators, such as IRSp53 and srGAP2. IRSp53, a member of the I-BAR family found in lamellipodia and filopodia has been shown to bind Cdc42, Rac1 and WAVE2^71,72^. Other known GAPs, such as ARHGAP22, ARHGAP24 (FILGAP), and SH3BP1 interact with the proteins involved in cell protrusion and could play a similar role as β2-chimaerin. In particular, it was previously shown that depletion of SH3BP1 resulted in a high activity of Rac1 at the front^73^. Thus, even if β2-chimaerin was sufficient to explain Rac1 shaping in the present work, we cannot exclude that other Rac1 GAPs also play a role, through redundant activities or additional levels of regulation. Further work should be performed to screen exhaustively the contribution of each of these GAPs. Yet, in the context of our findings, we can expect that one or a combination of GEFs for Rac1 and Cdc42 should form effective gradients of ~10μm decay length that would act as the primary spatial cue for these Rho GTPases. It remains to be explored if the Cdc42 and Rac1 positive feedbacks and crosstalks, as previously suggested^9^ and observed in our work (**Fig 3b**), play a role in shaping GEF distributions.

## Materials and Methods

### Plasmids and molecular constructs

ITSN-CRY2-mCherry was constructed as detailed previously^49^. The TIAM DH-PH domain was similarly amplified from TIAM(DHPH)-Linker-YFP-PIF (gift from O. Weiner, University of California, San Francisco) and cloned into CRY2PHR-mCherry. Both ITSN-CRY2-mCherry and TIAM-CRY2-mCherry were cloned in pHR lentiviral vectors (gift from O. Weiner) using MluI and BstBI cloning sites. N-WASP-iRFP, PAK1-iRFP, Abi1-iRFP and β2-chimaerin-iRFP fusion genes were constructed by cloning the corresponding human cDNAs upstream the iRFP713 gene sequence^74^, separated by a PVAT sequencer. Rac1BS and Cdc42BS plasmids were kindly provided by Dr Louis Hodgson^40^, and were subcloned into the lentiviral pLVX vector (Clontech, Mountain View, CA USA) between XmaI and XbaI cloning sites.

### Cell culture and reagents

HeLa cells were cultured at 37°C with 5% CO2 in DMEM (Dulbecco’s modified Eagle’s medium) supplemented with 10% fetal bovine serum. Transfections were performed using X-tremeGENE 9 (Roche Applied Science, Penzburg, Bavaria, Germany) according to the manufacturer’s instructions using an equal amount of plasmid DNA for each construct (1 μg). Stable cell lines were obtained using lentiviral infections: all lentiviruses were produced by transfecting pHR- or pLVX-based plasmids along with the vectors encoding packaging proteins (pMD2.G and psPax2) using HEK-293T cells. Viral supernatants were collected 2 days after transfection and HeLa cells were transduced at an MOI of 2. Gene expression knockdown was achieved using pooled siRNA with the following sequences. Cdc42 : 5’- CGAUGGUGCUGUUGGUAAA-3’ and 5’-CUAUGCAGUCACAGUUAUG-3’, β2-chimaerin : 5’- AUUGAAGCAAGAGGAUUAA-3’ and 5’-CCACUUCAAUUAUGAGAAG-3’, ctrl : 5’- AGGUAGUGUAAUCGCCUUG-3’ and 5‘-GCGGGATATTTCGGTCAAT-3’. siRNA transfection was done following the manufacturer’s protocol (Lipofectamine RNAiMax, Thermo Fischer Scientific), and cells were imaged 48 hours after transfection. The JLY cocktail (8 μM jasplakinolide, 5 μM Latrunculin B, 20 μM Y27632) was applied 15 minutes before image acquisition.

### Live cell imaging and optogenetics

Micropatterned coverslips were prepared as described by Azioune et al.^75^. Releasable micropatterns were prepared similarly, with PLL-PEG being replaced by azido-PLL-g-PEG (APP) at 100 μg/mL. Migration was released by addition of 20 μM BCN-RGD for 10 minutes. Before imaging, cells were dissociated using Versene (Life Technologies) and seeded for adhesion on the previously mentioned coverslips for at least 2 hours. Experiments were performed at 37°C in 5%CO2 in a heating chamber (Pecon, Meyer Instruments, Houston, TX) placed on an inverted microscope model No. IX71 equipped with a 60X objective with NA 1.45 (Olympus, Melville, NY) and a Luca R camera (Andor, Belfast, UK). The microscope was controlled with the Metamorph software (Molecular Devices, Eugene, OR). TIRF images were acquired using an azimuthal TIRF module (iLas2; Roper Scientific, Tucson, AZ). Optogenetics stimulations were performed every 30-40s with a DMD (DLP Light Crafter, Texas Instruments) illuminated with a SPECTRA Light Engine (Lumencor, Beaverton, OR USA) at 440 ± 10 nm.

### FRET

HeLa cells were lentivirus-infected with a Cdc42-FRET-biosensor or a Rac1-FRET-biosensor (kindly provided by Louis Hodgson) and FACS sorted for intermediary fluorescence. 24 hours after plating them on glass coverslips, cells were imaged by TIRF microscopy. Excitation was done with a laser at 405 nm, dichroic mirrors stayed the same (BS: FF-458-DiO2, Semrock) while a filterwheel allowed for the switching of appropriate emission filters to acquire sequentially donor (mCerulean, Em : FF01-483/32) and FRET (Em : FF01-542/27) emissions. Image processing included registration, flat-field correction, background subtraction, segmentation and FRET/donor ratio calculations. FRET profiles measured from the FRET images were normalized between 0 and 1 when comparing the two Cdc42 and Rac1 FRET reporters or when comparing a FRET reporter with another fluorescence signal. We did not normalize FRET profiles from the same reporter when comparing two different experimental conditions.

### Immunofluorescence Microscopy

HeLa cells stably expressing a Rac1-FRET-biosensor were fixed in PBS with 4% PFA for 20 minutes at 25°C. After permeabilization in PBS + 0.1% triton X-100 for 15 minutes and blocking in PBS + 1% BSA + 1% FBS for 20 minutes, stainings were performed in PBS with 0.05% triton + 1% BSA 1hr at room temperature. Antibodies were used as follows: β2-chimaerin primary antibody: 1 /100 (Orb182594, Biorbyt), anti-rabbit-Alexa594 antibody: 1 /400 (ThermoFisher). Acquisitions were made in HiLo mode using an azimuthal TIRF module as described above).

### Image processing and quantification of intracellular gradients

Images were analyzed with custom-built Matlab routines. For the images obtained in our optogenetic experiments, we subtracted the initial pre-optogenetics signal from all subsequent images in order to measure solely the recruitment of fluorescent proteins to the basal membrane and to avoid volume artifacts. The resulting differential images where normalized between 0 and 1 (zero being the background value outside cells) and then averaged over 10 time points and over all cells to produce the averaged images shown in **figures 2d,f** and **3b,d** (displayed with the “fire” color scale). For the quantification of the gradients presented in **figure 1b**, and **3c,f,g,** we measured the FRET ratio along 2 linescans per cell, drawn manually perpendicular to the cell edge in protrusive regions with a line width of 10 pixels. The gradients were first averaged for each cell, and then averaged over all cells. For the quantification of the gradients presented in **figure 2b,e,g** and **3e,h,i**, fluorescence was quantified along a line of 10 pixels in width spanning across the cell diameter in the direction of the optogenetic gradients. The curves in **Fig 2b,e,g** and **3e,h,i** were normalized between 0 and 1 where 0 stands for the average of the 5 minimal values and 1 stands for the average of the 5 maximal fluorescence values.

### Processing of the migration movies

Movies were analyzed with custom-built Matlab routines. The segmentation of cell borders was performed on fluorescence images using the Matlab function “Graythresh”. Cell centroid positions were determined using the Matlab function “Regionprops” and used to quantify cell movement. To measure cell velocity, we computed instantaneous speed of cell centroids at each time frame, and then averaged it over several time frames. Cells stimulated through TIAM activation reached maximum speed soon after the beginning of illumination, so instantaneous speed was averaged between t=15 minutes to t=45 minutes. Cells stimulated through ITSN activation reached maximum speed at later stages, and instantaneous speed was thus averaged between t=60 minutes to t=90 minutes. Angular precision was computed as follows: for each cell, the displacement vector was computed between the initial cell centroid (averaged over the 3 first time frames) and the final cell centroid (averaged over the 3 last time frames), and we measured the angle between this vector and the axis of stimulation gradients. These angles were bootstrapped over 1000 replications, and angular precision was estimated with the formula 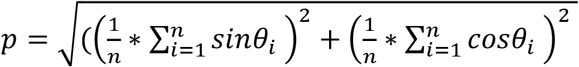 morphodynamics maps (**Fig 5c**) were obtained using a routine adapted from (Yang 2015). The cell contour was aligned such that the middle of the map was centered on the direction imposed by the optogenetic gradient.

### Data and code availability statement

The data that support the findings of this study and all custom codes used for analysis are available from the corresponding author upon reasonable request.

## Acknowledgments

We thank Louis Hodgson for sharing the Rho GTPase sensors; Maria Carla Parrini, Kristine Schauer and Christophe Tribet for critical reading of the manuscript and helpful discussions; Remy Fert and Eric Nicolau for their technical assistance; The Institut Curie Shared FACS Facility for cell sorting. M.C. acknowledges financial support from French National Research Agency (LICOP n° ANR-12-JSV5-0002-01). M.C. and M.D. acknowledge funding from French National Research Agency (ANR) Paris-Science-Lettres Program (ANR-10-IDEX-0001-02 PSL), Labex CelTisPhyBio (N° ANR-10-LBX-0038), the France-BioImaging infrastructure supported by ANR Grant ANR-10-INSB-04 (Investments for the Future), and Institut Pierre-Gilles de Gennes (laboratoire d’excellence, “Investissements d’avenir” program ANR-10-IDEX-0001-02 PSL and ANR-10-LABX-31).

## Author Contributions

Conceptualization, S.d.B., M.D., and M.C.; Methodology, S.d.B., M.D. and M.C.; Software, S.d.B. and M.C.; Formal Analysis, S.d.B., K.V. and M.C.; Investigation S.d.B., K.V. and J.M.; Resources, F.D.F., G.C., and F.D.; Writing – Original Draft, S.d.B. and M.C.; Writing – Review & Editing, S.d.B., M.C., M.D. and K.V.; Visualization, S.d.B. and K.V.; Supervision, M.C.; Funding Acquisition, M.C.

